# A Novel Work Flow for Health Survey Analysis: Results on the Yazd Health Study Data (YaHS)

**DOI:** 10.1101/491142

**Authors:** Elyas Heidari, Vahid Balazadeh-Meresht, Nastaran Ahmadi, Mahmoud Sadr, Ali Sharifi-Zarchi, Masoud Mirzaei

## Abstract

Health surveys are one of the most prominent sources of information in health and medical sciences. Analysis of a health survey not only gives information about the inherent features of the population under study but also sheds light on the interactions between different health factors. Even though there has been a multitude of methodologies proposed for survey analysis, many of them are biased towards the previous hypotheses, the fact which may lead to spurious results by neglecting the whole structure of relations among the factors. Here, we proposed a novel work flow on survey analysis. By gathering the previously less-utilized graphical models on survey analysis, newly-proposed methods, and commonly-used tools of prediction and hypothesis testing our work flow provides the medical community with an end-to-end pipeline for comprehensive statistical analysis of health surveys. To demonstrate the functionality of our work flow, we validate it on the Yazd Health Study (YaHS) dataset, in which a questionnaire of 300 questions from 10000 participants associated with 40 laboratory measurements from a subset of 4010 individuals is acquired from the residents of Yazd Greater Area of Iran. Many of our findings are aligned with previous medical knowledge. Most of our new findings are surprisingly observed in the clinics, each of which is potential to raise a promising hypothesis.

## Introduction

Including measurements of demographics, health status, nutrition habits, habitual behaviors, socioeconomic status, etc., health surveys can be cited as the chief informative source about public health. Merits of analyzing health surveys are manifold. Eliciting information about the outcome of the provided public health services form health surveys, health care policymakers can regulate their policies of health-related programs. The importance of health surveys is even more evident in the societies which are less studied. By so doing, information and knowledge about rarely known populations can be made and appended, in turn, to the current knowledge of global health, thereby, elucidate new homogeneity and heterogeneity of health regimes among the global population. From another perspective, analysis of health surveys, in addition to uncovering several less-/unknown statistical relations among different measurements, can be used to shed light on some less investigated measurements (variables) which are potential to be instructive to medicals, specifically in designing new surveys. Furthermore, health surveys are essential when someone is looking for the risk factors of a specific disease or crosswalk of different conditions, regarding the aforementioned measurements.

To date, quite a few approaches have been put forward for analyzing health surveys [1][2][3][4][5]. Regression analysis, correlation analysis, causal analysis, and hypothesis testing are among the most commonly-used analyses on survey data [6][7][8][9]. Although all of these approaches made remarkable findings, many of them have been biased in favor of some predetermined investigations (e.g., particular associations among variables), in light of the underlying complexity of the health surveys. In analyzing a health survey, one should take different data types (i.e., discrete, categorical, or continuous) into account. Thus, they should utilize methods which apply to different types of data variables. Furthermore, considering numerous variables intensifying the intricacy of the analysis, it is indispensable to employ methods streamlining the analysis as well as interpretation in such high-dimensional settings.

Yazd Health Study (YaHS) is a population-based prospective study of 10000 Yazd adult residents aged 20-69 years, conducted in November 2014 in Yazd Greater Area of Iran [10]. A verified questionnaire including 300 questions regarding a) demographics, b) physical activity, c) sleep quality and quantity, d) mental health, e) past medical history of chronic disease and surgical operations, f) dental health, g) history of accidents, h) dietary habits, i) occupation and social life, j) traditional medicine, k) smoking habit and drug addiction, l) women’s health and m) quality of life is recorded by trained interviewers. Moreover, anthropometrics, blood pressure, and vital signs are measured from a subset of 4010 who participated in blood testing.

Here, we have 37 continuous variables and 172 categorical variables after preprocessing the data. Hence, unbiased inspection of all of the associations among variables is not practically feasible with the approaches mentioned above. We exploit methods which give us an estimation of the whole structure of the relations to be subsequently used for investigating a smaller set of possible relations. To this end, we use probabilistic graphical models (PGMs) which accordingly estimate a sparse structure of relations among variables. The estimated structures are then used for statistical assessment of the relations. PGM facilitate an unbiased analysis in multivariate settings. Graphical models have been used previously in various fields of research, including bioinformatics [11][12][13][14][15][16][17][18], social and internet sciences [19][19][20][21][22], and economics [23][24][25][26]. In this analysis we use three types of graphical models: i) Gaussian graphical models, for associations among continuous variables; ii) minimal forest, for associations among the whole set of variables, including both categorical and continuous variables; and iii) causal network, for causal relations among continuous variables. Our analysis with PGMs is analogous to widely-used correlation analysis, but there are few differences (e.g., transitivity of regular correlation). After that, we conduct exploratory data analysis (e.g., with scatter plots, box plots, or heat maps) to assess the findings from PGMs, more accurately.

In addition to estimating the structure of associations among variables, some other questions are appealing to be answered. One of these questions is that what are the differences between the two separated groups of samples, for instance, between a group of patients and a group of healthy participants. In other words, which variables, including laboratory measurements, lifestyle, or familial health profile are significantly different between two distinct groups of samples. As a solution to this question, some of us has recently proposed variable-wise Kullback-Leibler divergence (VKL) [27]. Relations between various pairs of health factors have been suggested abundantly in the literature. Associations between fat and fasting serum glucose (FSG), fat and blood pressure, and body mass index (BMI) and blood pressure have been proposed frequently [28][29]. However, in each experiment, there are some samples which do not follow such expected relations. For instance, there are some individuals with hypertension and low BMI or low FSG and high bodily fat. Therefore, it is admissible to suppose that such samples bear some subtle features violating or blocking the aforementioned expected relations. To discover the violating features, we use a recently proposed method, violating variable-wise Kullback-Leibler divergence (VVKL) [27].

As mentioned before, a prevalent analysis for surveys is to regress one variable on other variables. This analysis in addition to estimating weights of exposure variables in predicting the response variable can be generalized by regularization (penalization) for selecting active variables and excluding confounding variables in predicting the response variable. Hereafter, we also use the elastic net model to predict values of an objective variable based on all other variables. Elastic net selects a scars number of variables to predict the variable of interest thereby proposing a few variables to be associated with the response variable, similar to the PGM and VKL.

In this article besides proposing a new end-to-end pipeline for statistical analysis and interpretation of health surveys, we report some of our noteworthy findings of the analysis of the YaHS dataset. Alignment of the majority of non-trivial findings in this analysis with the previous knowledge boosts the reliability of our new findings. Such results which are underscored by several statistical methods performed in this analysis can be tested as plausible hypotheses by medical researchers.

## Results

### 2.1 Structure of the population

To capture the structure of the population, we reduce the dimension of the data from 37 (number of laboratory measurements) to 2 with Uniform Manifold Approximation and Projection (UMAP). As indicated in fig. 1a samples with different genders and age groups are dispersed throughout the population. This demonstrates that our population is fairly homogeneous concerning the laboratory measurements. To delineate the overall structure of the population, some statistics are shown in fig. 1b to fig. 1c.

**Figure 1:**
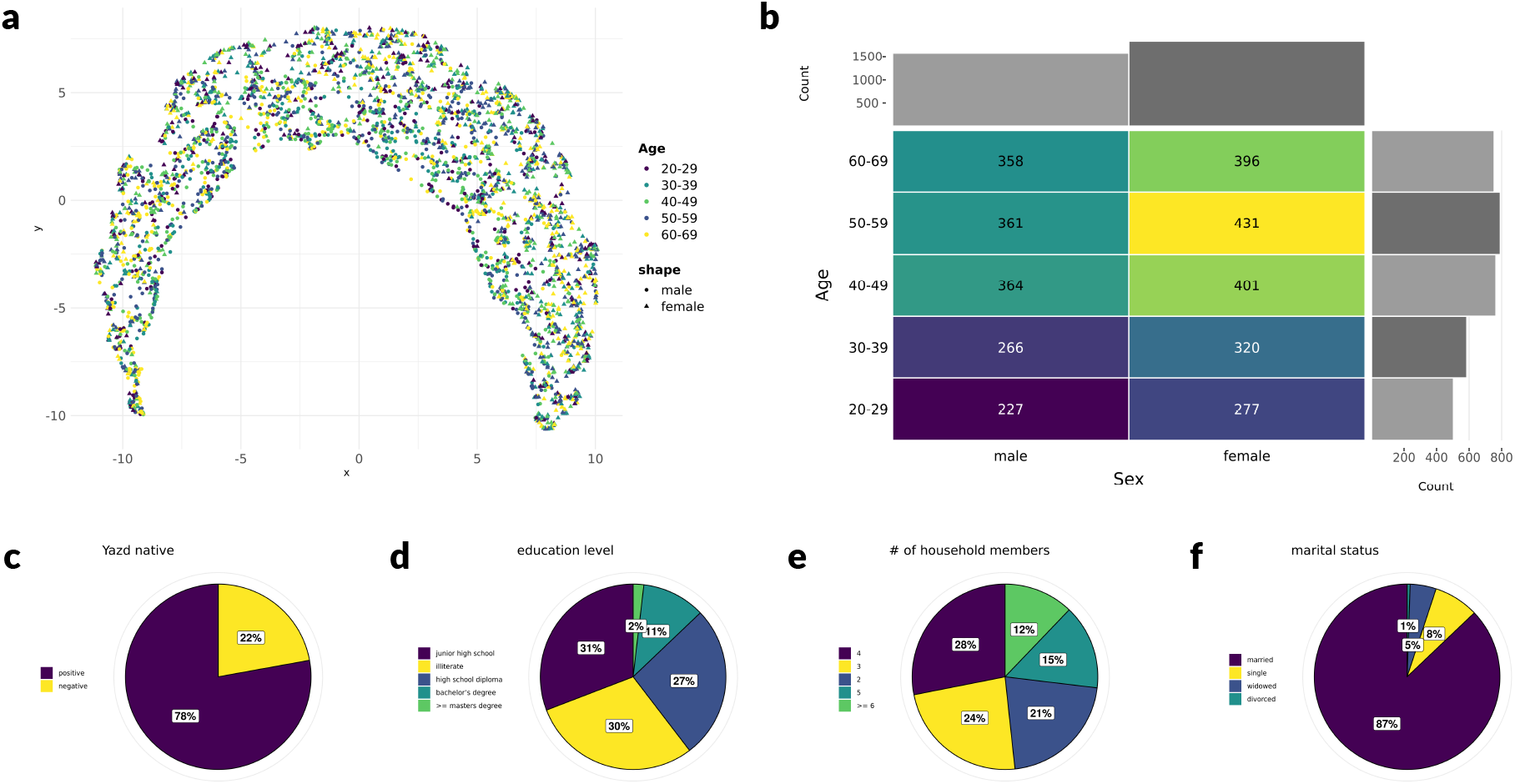
Overview of the population. **(a) UMAP plot of the population:** The population is plotted in 2 dimensions with UMAP, each point represents a sample from the population. Points are colored with respect to their age group and their shapes indicates the gender of the corresponding sample. **(b) Distribution of sex and age:** Each cell represents the number of samples with specific sex and age group. The bar plots on the top and the right show distribution of sex and age, respectively. **(c to f) Pie charts of 4 measurements: (c)** nativity of Yazd, **(d)** education level, **(e)** number of household members, **(f)** Martial status.

### 2.2 Gaussian graphical model of laboratory measurements

We employ graphical models to capture a sparse network of interactions (relations) among the variables. A Gaussian graphical model (GGM) is used to construct the graph of associations among continuous variables in which each node represents a variable, and each edge demonstrates a non-zero partial correlation between to variables. We use a community detection algorithm to find the communities of highly related variables within the graph. As illustrated in fig. 2a, the GGM is decomposed into five communities, where the communities 1 and 2 indicate metabolic measurements and the communities 3 and 4 indicate immunological and hematological measurements. To investigate the strength of the relations, we calculate the mutual information between each pair of the variables fig. 2b. Moreover, the summation of the mutual information with all other variables is computed for each variable as a criterion for the informativeness the variable. A summary description of the variables is reported in 1. The results emphasize the importance of neck circumference as the variable with the most degree in the GGM and among the most important variables regarding betweenness centrality and mutual information. Furthermore, hemoglobin has the most betweenness centrality in the graph and links the blood cell communities (3 & 4) to the metabolic communities (1 & 2). Creatinine is also highlighted with the most cumulative mutual information value. While most of the relations are consistent with the previous results in the literature, interestingly, some links are less investigated/proposed.

**Figure 2:**
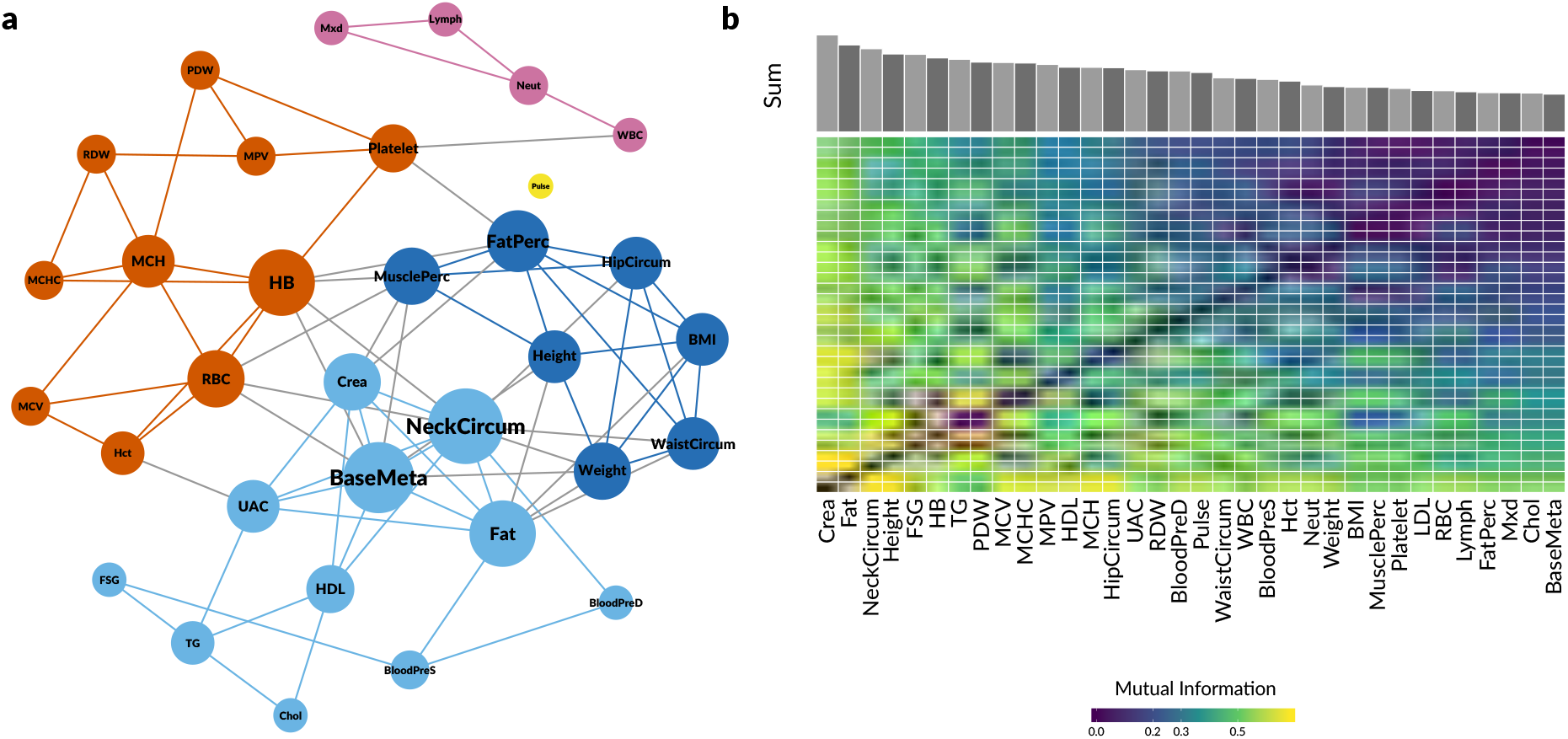
Overview of the relations among the continuous variables. **(a) Structure of the GGM:** Nodes represent continuous variables (laboratory measurements) and edges indicate non-zero partial correlations between pairs of variables. Nodes are colored based on their node community, and their sizes show their betweenness centrality in the graph. There are five communities in the graph each of which consist of variables with a class of functionality. Communities 1 and 2 are about metabolic indices, community three is about hematologic cell counts, and community 4 includes white cell counts. **(b) Heat map of mutual information among the variables:** Each cell indicates the mutual information between two continuous variables. The summation of the mutual information for the variables are indicated in a bar plot on the top.

**Figure 3:**
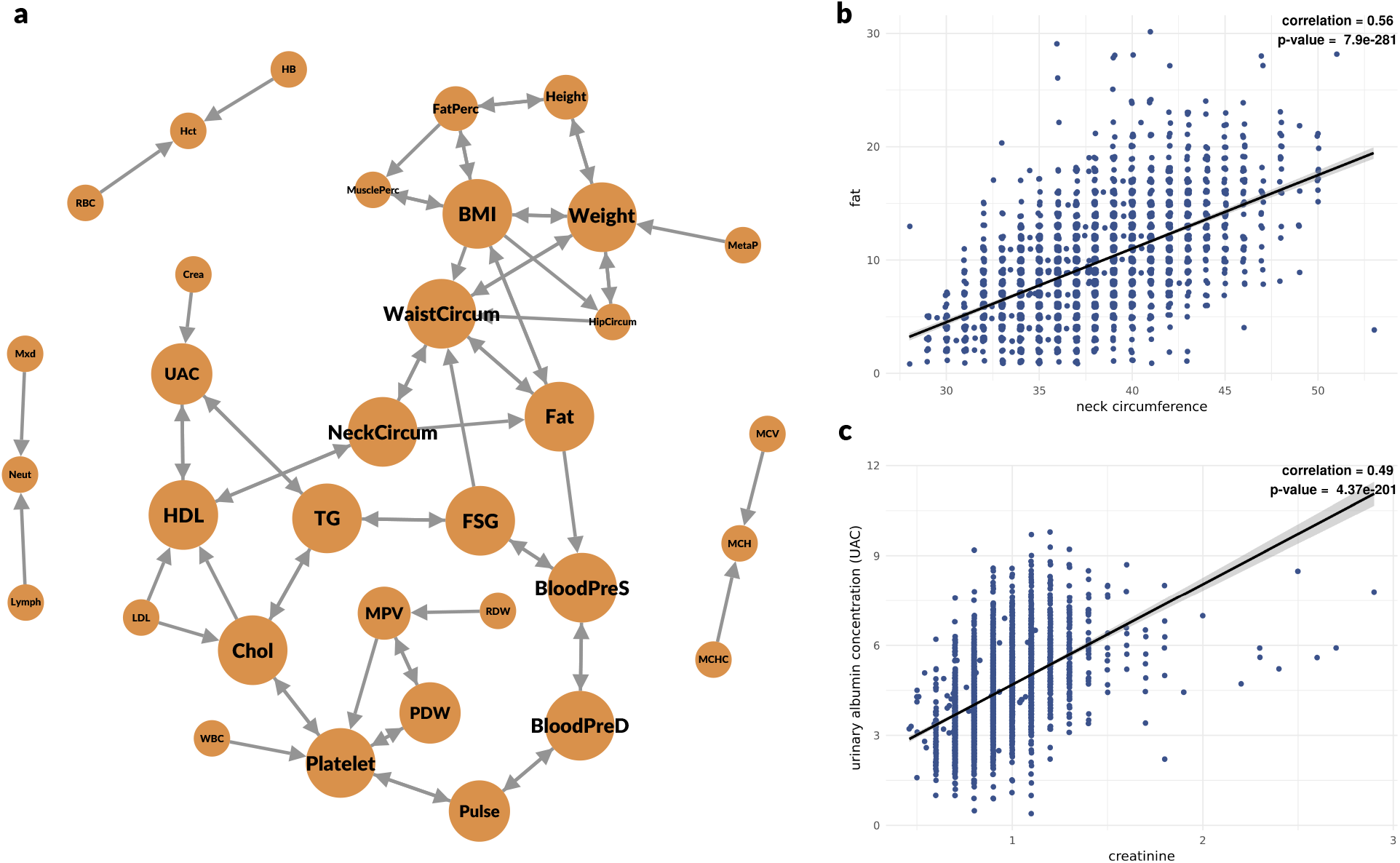
Results from the causal model. **(a) Structure of the causal network:** Nodes represent continuous variables (laboratory measurements) and edges indicate causal relations between a pairs of variables. Each edge is directed from cause to effect and bi-directed edges either indicate a two-way causal relation or an uncertainty in the direction of the causal relation estimated by the causal model. **(b & c) Scatter plots of the associations extracted from the causal network:** Each blue point represents a sample, Pearson correlation and p-value of correlation test for the association are reported on the top-right position of each scatter plot. **(b)** neck circumference and fat, **(c)** creatinine and UAC.

### 2.3 Minimal forest of all of the variables

We exploit the minimal forest algorithm for estimating relations among a mixed set of variables including both continuous and categorical variables. Again, each node represents a variable, and each edge represents a statistical association between two nodes. After constructing the graph, we use a community detection algorithm to find the communities of highly related variables. To analyze the network, we explore each community focusing on its variables. The structure of the minimal forest is illustrated in fig. 4 and some findings are depicted in fig. 5. The communities are analyzed in detail in the following.

**Figure 4:**
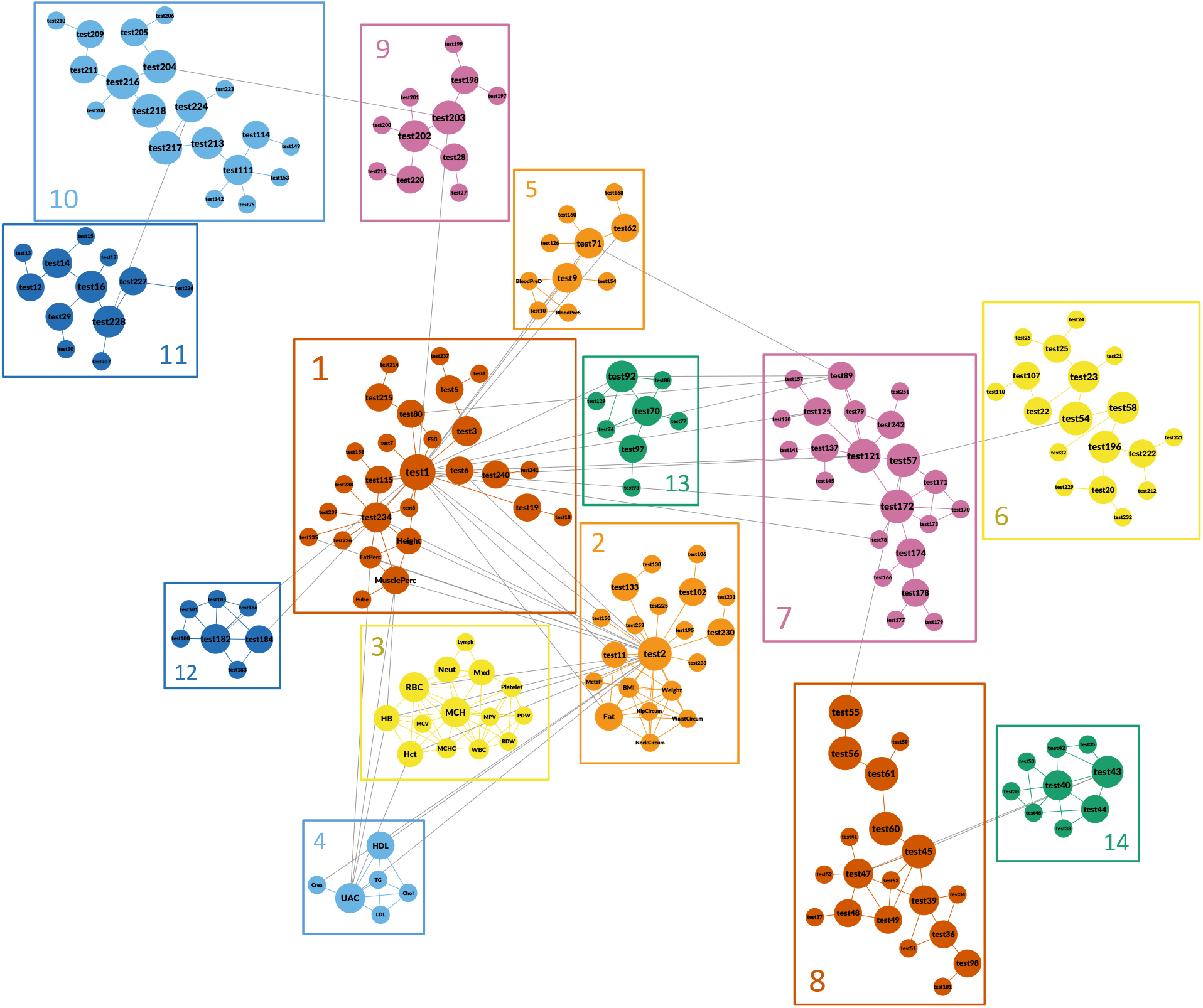
Structure of the minimal forest. Nodes represent variables and edges indicate estimated relations based on mutual information between pairs of variables. Nodes are colored based on their node community and their sizes show their betweenness centrality in the graph. There are 14 communities in the graph each of which consist of variables with a class of functionality: **1)** age overall health profile; **2)** gender, metabolic variables, etc.; **3)** blood cell counts; **4)** measure biochemistry features; **5)** blood pressure and heart disease; **6)** sleeping and health status assessment; **7)** bodily pain, osteoporosis, referring to doctor or hospital; **8)** bodily and mentally well-being; **9)** nutrition habits and TV usage; **10)** nutrition habits, AD, etc.; **11)** diary usage and physical activity; **12)** dental health; **13)** familial history of some diseases; **14)** mental health.

**Figure 5:**
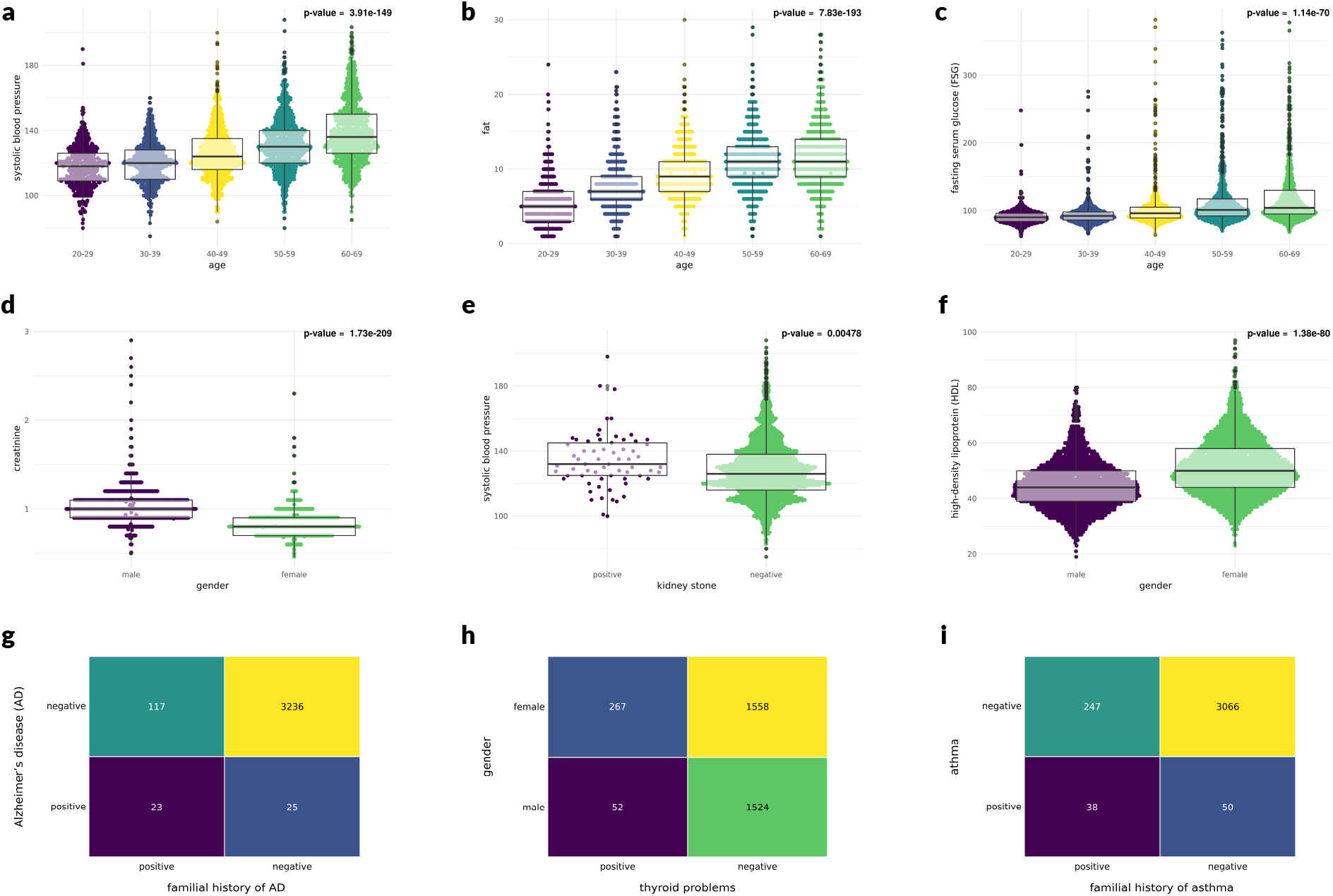
Results from the minimal forest. (**a to f**) **Box plots of the associations extracted from the minimal forest:** Each point represents a sample, p-value of ANOVA for the association are reported on the top-right position of each box plot. **(a)** Age and systolic blood pressure, **(b)** age and fat, **(c)** age and FSG, **(d)** gender and creatinine, **(e)** kidney stone and systolic blood pressure, **(f)** gender and HDL. **(g to i) heat maps of the associations extracted from the minimal forest:** each cell represents the number of samples with specific values of the two variables in the heat map. **(g)** familial history of AD and AD, **(h)** thyroid problems and gender, **(i)** familial history of asthma and asthma.

- **Community 1** is the central community of the network, and test 1 (age) is the center of this community. Test 8 (marital status) is connected to test 1 (age), this node is connected to test 234 (occupation status), and tests 236, 239, and 238 are connected to test 234 and are also about the occupation. Test 234 is connected to test 2 (gender) demonstrating the difference of occupation status between males and females. Test 19 (get-up time) from this community is connected to test 1. fasting serum glucose (FSG) is also connected to test 1. Tests 3, 4, and five which are about residential status are in the community 1 and are connected to test 1 through test 3. test 6 (education status) is also connected to test 1. Test 1 is additionally connected to test 9 (blood pressure), from the same community, and test62 (heart disease), form community 5; fat and body mass index (BMI, test 11) from community 2; and community 12 which indicates dental health. All in all, community 1, as a neighborhood of age, describes the general health status.
- **Community 2** is centered at test 2 (gender). Test 2 is connected to a highly connected group of nodes, including fat, BMI, MetaP, and body circumferences. Also, this node is connected to some nodes from community five which represents the red blood cell counts. Test 2 is also connected to community 5 through (high-density lipoprotein) HDL and creatinine. Test 102 (thyroid problems) is also connected to test 2.
- **Community 3** consists of hematological and immunological measurements.
- **Community 4** includes urinary albumin concentration (UAC), creatinine, cholesterol, and triglyceride. All of these variables are from biochemistry profile.
- **Community 5** encompasses nodes about blood pressure, hypertension, heart disease, skin cancer, ovarian cancer, and kidney stone. Intriguingly, tests about cancer and kidney stone are related to blood pressure and hypertension. Furthermore, there is an edge between test 71 (hypertension) and test 89 (high blood cholesterol).
- **Community 6** is connected to the rest of the network through the edge between test 54 (health status assessment) and test 57 (bodily pain). Test 54 is connected to test 23 (not fall asleep within 30 minutes) and test 58 from the same community. Test 23 is connected to the other questions about sleep quality. Test 22 (using sleeping pills) is connected to test 107 (depression). Test 107 is connected to test 110 (familial history of depression). Test 58 (bodily energy) is connected to test 32 (number of close friends). From this community, the relation of test 20 (duration of sleep) with test 232 (putting a salt pot on the table) and test 229 (diary usage) is also appealing.
- **Communities 7 and 8** include all of the tests about mental health and are connected through the edge between test 55 and test 57 to the rest of the graph. Test 98 (asthma) is also in this community and is connected to test 36 (trouble in breathing) and test 101 (familial history of asthma).
- **Communities 9, 10, and 11** contain all of the variables about nutrition, TV usage, internet usage, and physical activity. The blue community also includes test 111 (Alzheimer’s disease). Test 111 is connected to test 75 (history of mental illness) and test 114 (familial history of Alzheimer’s disease).

### 2.4 Causal Inference

To find the causal interactions between continuous variables, we use a Bayesian Network in which each node represents a variable, and each edge is a causal relation with its arrow indicating the direction of the causal relation. After construction of the graph, we use a community detection algorithm to find the communities of highly related variables within the graph 3.

### 2.5 Variable-wise Kullback-Leibler divergence (VKL)

Variables which their distribution differ significantly between two specific groups of samples are informative, given the fact that they can be recognized as causes, effects, or both cause and effect of a particular feature. For instance, the variables which their distributions are exceptionally different between two groups of normal samples and patients can be remarked as disease hallmarks. To detect such variables for different diseases we employ VKL which exploits KL divergence to find the variables which differ the most between two groups of normal and affected samples. We apply VKL separately on the laboratory measurements and the questions (tests) for heart disease (fig. 6), thyroid problems (fig. 7), and osteoporosis (fig. 8). In the following subsections, we will report our findings in the VKL analyses.

**Figure 6:**
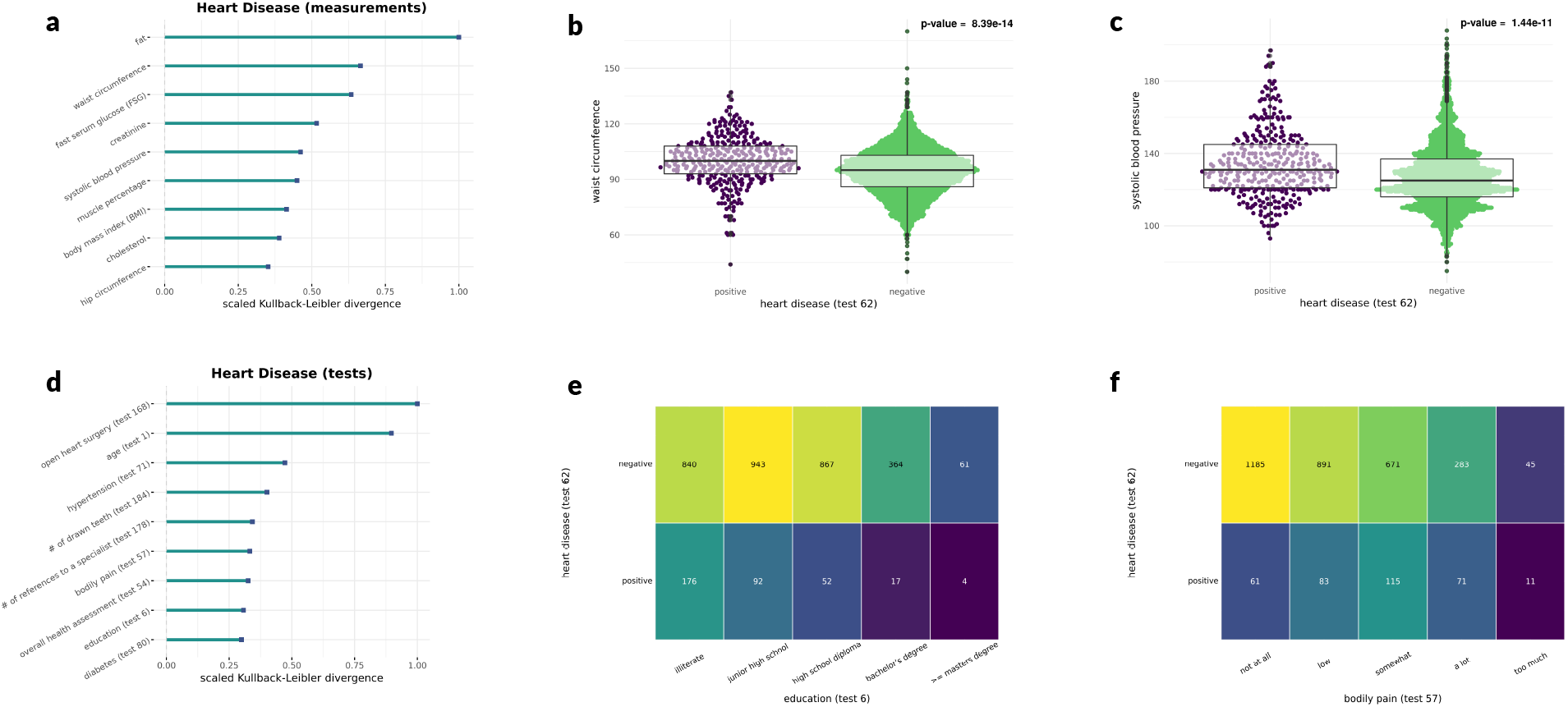
VKL analysis for heart disease. **(a) VKL bar plot:** After excluding trivial variables, we selected continuous variables with the highest KL values, each bar shows the scaled KL value. KL values are scaled such that each KL value is divided by the highest value among all KL values. **(b & c) Box plots:** Two arbitrary continuous variables detected by VKL are plotted to show the differences between two groups of people with and without heart disease. **(b)** Waist circumference, **(c)** systolic blood pressure. **(d) VKL bar plot:** same as (a) but for categorical variables. **(e & f) Heat maps:** Two arbitrary categorical variables are plotted to indicated the differences between two groups of people with and without heart disease. **(e)** education, **(f)** bodily pain.

**Figure 7:**
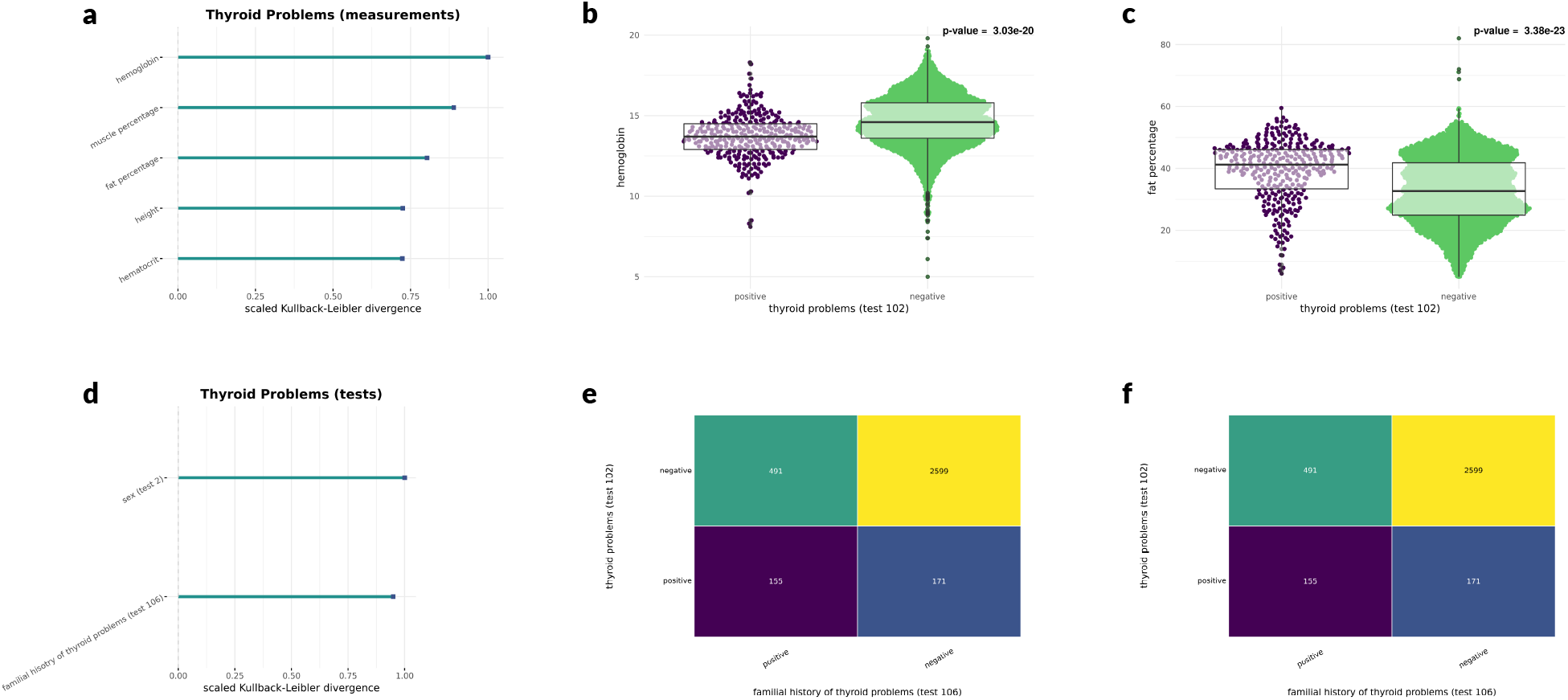
VKL analysis for thyroid problems. **(a) VKL bar plot:** After excluding trivial variables, we selected continuous variables with the highest KL values, each bar shows the scaled KL value. KL values are scaled such that each KL value is divided by the highest value among all KL values. **(b & c) Box plots:** Two arbitrary continuous variables detected by VKL are plotted to show the differences between two groups of people with and without thyroid problems. **(b)** Hemoglobin, **(c)** fat percentage. **(d) VKL bar plot:** same as (a) but for categorical variables. **(e & f) Heat maps:** Two categorical variables proposed by VKL are plotted to indicated the differences between two groups of people with and without thyroid problems. **(e)** Familial history of thyroid problems, **(f)** sex.

**Figure 8:**
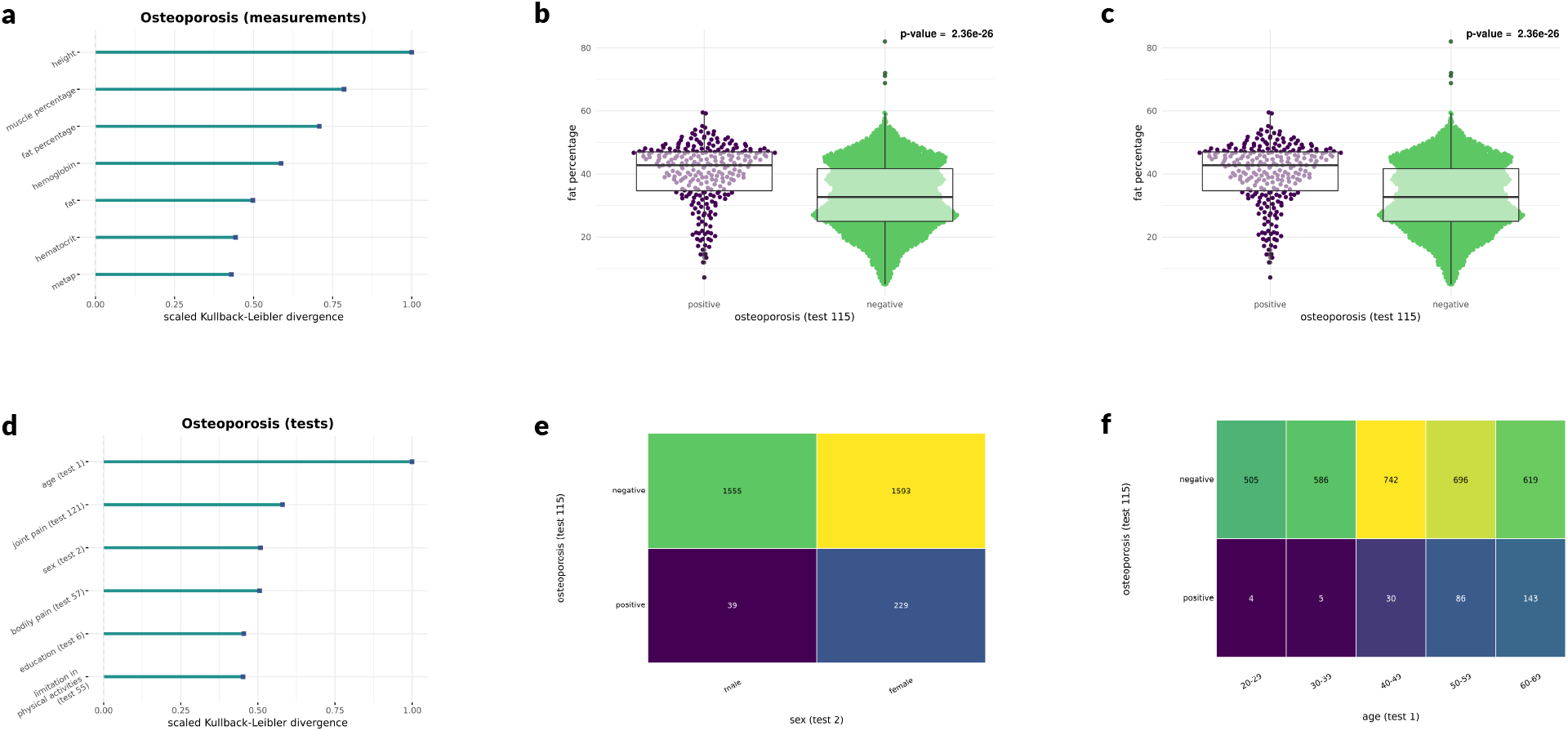
VKL analysis for osteoporosis. **(a) VKL bar plot:** After excluding trivial variables, we selected continuous variables with the highest KL values, each bar shows the scaled KL value. KL values are scaled such that each KL value is divided by the highest value among all KL values. **(b & c) Box plots:** Two arbitrary continuous variables suggested by VKL are plotted to show the differences between two groups of people with and without osteoporosis. **(b)** Hemoglobin, **(c)** fat percentage. **(d) VKL bar plot:** same as (a) but for categorical variables. **(e & f) Heat maps:** Two arbitrary categorical variables proposed by VKL are plotted to indicated the differences between two groups of people with and without osteoporosis. **(e)** Familial history of thyroid problems, **(f)** sex.

#### 2.5.1 Heart disease

As indicated in fig. 6a, VKL proposes fat, waist circumference, FSG, creatinine, systolic blood pressure, muscle percentage, BMI, cholesterol, and hip circumference to have most different distributions between groups of participants with and without heart disease, among laboratory measurements. Among these, waist circumference (fig. 6b) and systolic blood pressure (fig. 6c) is significantly associated with heart disease with p-values of 8.39e^−14^ and 1.44e^−11^, respectively (from ANOVA): Waist circumferences in participants suffering from heart disease are significantly higher. A similar pattern is observed for systolic blood pressure. Among questions (tests), experience of open heart surgery (test 168), age (test 1), hypertension (test71), number of drawn teeth (test 184), number of references to specialist in last year (test 178), bodily pain (test 57), overall health assessment (test 54), education level (test 6), diabetes (test 80) have the most different distributions between unaffected and affected groups of heart disease (6d). Although some variables like age, number of drawn teeth (which is highly correlated with age), and overall health assessment are trivial, interestingly, the distributions of bodily pain and education level show to be very different between affected and unaffected groups: people with heart disease suffer more from physical pain (6e), and more educated participants show to be less subjected to heart disease (6f).

#### 2.5.2 Thyroid problems

Hemoglobin, muscle percentage, fat percentage, height, and hematocrit are suggested by VKL to be highly associated with thyroid problems (fig. 7a). Specifically, hemoglobin (p-value = 3.03*e*^−20^) and fat percentage (p-value = 3.83*e*^-23^ are significantly different between two groups of participants with and without thyroid problems (fig. 7b & 7c): Hemoglobin levels in people with thyroid problems are lower than the group without thyroid problems. Same can be said about fat percentage. Also, sex (test 2) and familial history of thyroid problems (test 106) have different distributions within the groups above of samples (7d): people with the familial history of thyroid problems show higher tendency to be affected by thyroid problems (7e). Noteworthy, women tend to be more affected by thyroid problems (7f).

#### 2.5.3 Osteoporosis

As VKL asserts, height, muscle percentage, fat percentage, hemoglobin, fat, hematocrit, and metap are variables with the most different distributions between two groups of osteoporosis-affected and - unaffected individuals (8a). Remarkably, fat percentage (p-value = 2.36*e*^−26^) and age (p-value = ) are significantly related to osteoporosis (8b & 8c). Furthermore, age (test 1), joint pain (test 121), sex (test 2), bodily pain (test 57), education level (test 6), and limitation in physical activity (test 55) are reported by VKL (8d). Notably, females are more prone to be affected by osteoporosis (8e). Also, osteoporosis is more abundant in older people (8f).

### 2.6 Violating Variable-wise Kullback-Leibler divergence (VVKL)

Although associations between pairs of variables are well-known, such associations are either trivial (e.g., the association between hemoglobin and red blood cell count) or proved (e.g., the association between fat and blood pressure), in each experiment, there are some unexpected observations. For instance, in each health experiment in which blood pressure and fat are measured, there are some participants with relative low/high blood pressures given their fat profiles. Such unexpected observations can be assumed to bear some features which violate (or block) the expected associations. Seeking to find such violating features in unexpected observations we use VVKL. We applied VVKL to two renowned associations between fat and blood pressure (fig. 11) and between BMI and blood pressure (??). Details of the findings in the VVKL analyses come in the following.

#### 2.6.1 VVKL analysis of fat and blood pressure

Fat and blood pressure are known to have an intimate association; our data also support this fact (fig. 11a). Nevertheless, there are some samples with relatively high (green color in fig. 11a) or low (purple color in (fig. 11a)) blood pressures with respected to their bodily fat levels. Such groups are presumably bearing some features violating the expected linear association between fat and blood pressure. We, in turn, use VKL method to find the variables with differ the most between these two groups. Variables suggested by VKL can be remarked as the aforementioned violating features. Age (test 1), education (test 6), time spent on electronic messengers (test 241), FSG, amount of fast food usage per months(test 202), and number of drawn teeth (test 184) are the variables with the most different distributions between the two groups of violated samples (fig. ??b). Markedly, FSG show be much

### 2.7 Multivariate elastic net models

We fitted a multivariate elastic net model to predict the laboratory measurements and the state of bearing diseases concerning the laboratory measurements. The elastic net induces a sparse selection of the predictors for the model and, therefore, selects a subset of the variables for the prediction which is of high interest in medical research. Some of the results indicated in fig. 9 and fig. 10, demonstrate how precisely such models introduce new formulas for the health indices and candidate biomarkers of the diseases. The trend of changes of the predictors regarding the response variable is evident in fig. 9 and fig. 10.

**Figure 9:**
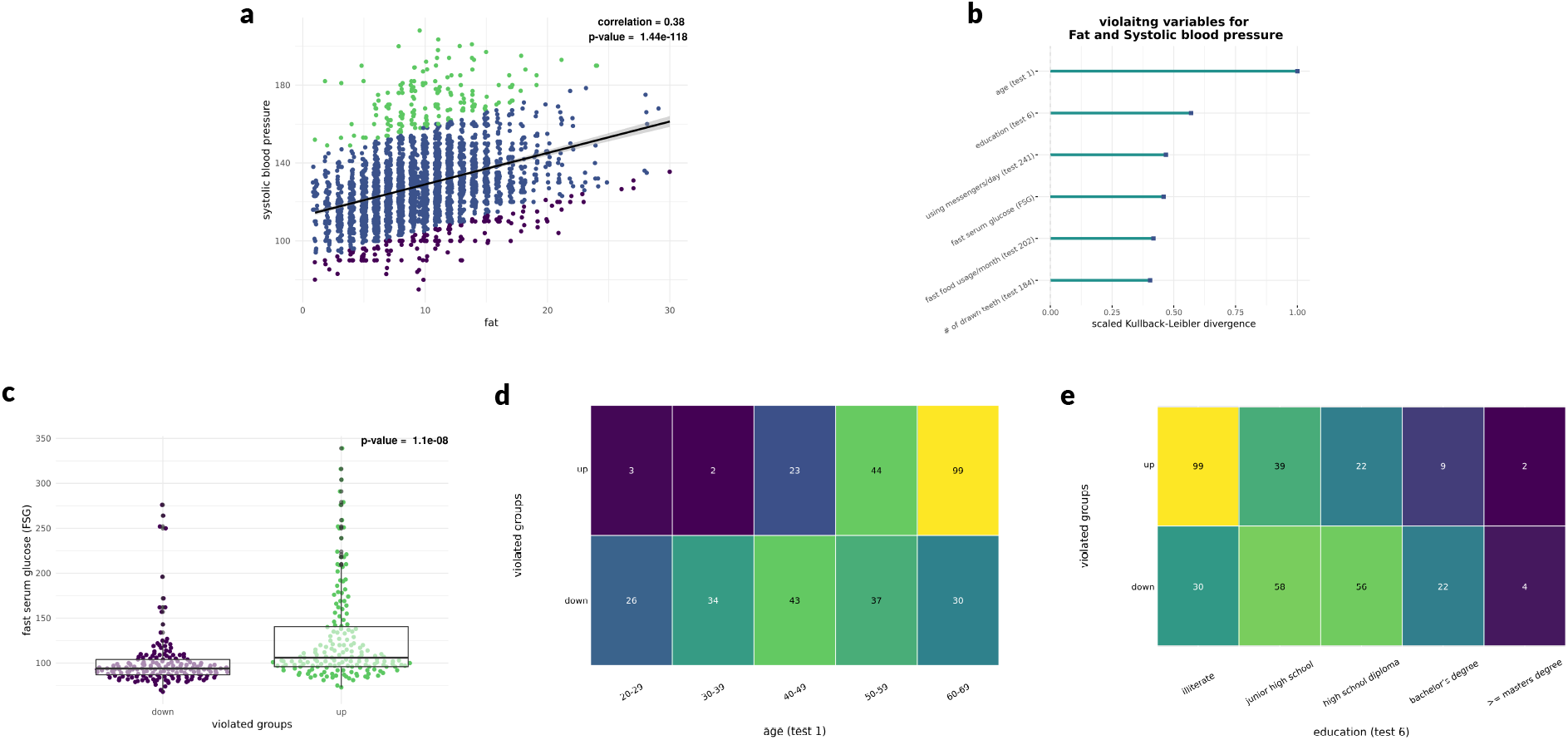
VVKL analysis for the association between fat and systolic blood pressure. **(a) Scatter plot for fat and systolic blood pressure:** Each blue point represents a sample, Pearson correlation and p-value of correlation test for the association are reported on the top-right position of the scatter plot. The violated groups are colored in green (up) and purple (down). **(b) VKL bar plot:** After excluding trivial variables, we selected continuous variables with the highest KL values, each bar shows the scaled KL value. KL values are scaled such that each KL value is divided by the highest value among all KL values. Each variable is supposed to violate the expected linear relation between fat and systolic blood pressure. **(d) Box plot of FSG concerning two violated groups of samples:** Each point represents a sample, the p-value of ANOVA for the association are reported on the top-right position of the box plot. **(d & c) Heat maps concerning two violated groups of samples:** each cell represents the number of samples with specific values of the variable within two violated groups of samples. **(d)** age, **(e)** education.

**Figure 10:**
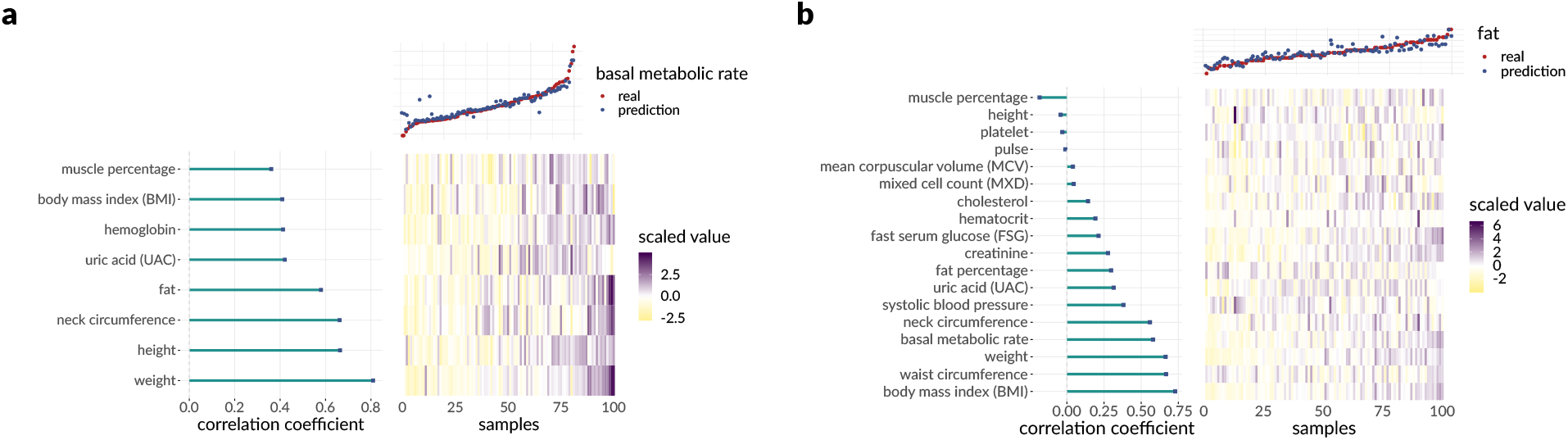
Multivariate elastic net model for continuous variables. **(a-b) An overview of the prediction model:** The predictors with a non-zero coefficient of in the model are taken into account. The scaled values of the predictors for a subsample of 100 participants are shown in the heat map. The correlation coefficients of the predictors and the response variable are shown as a horizontal bar plot on the left. A scatter plot depicting the real values (red dots) as well as the predicted values (blue dots) of the response variable is illustrated on the top, where the samples are sorted concerning the value of the response variable. **(a)** Basal metabolic rate, **(b)** bodily fat.

**Figure 11:**
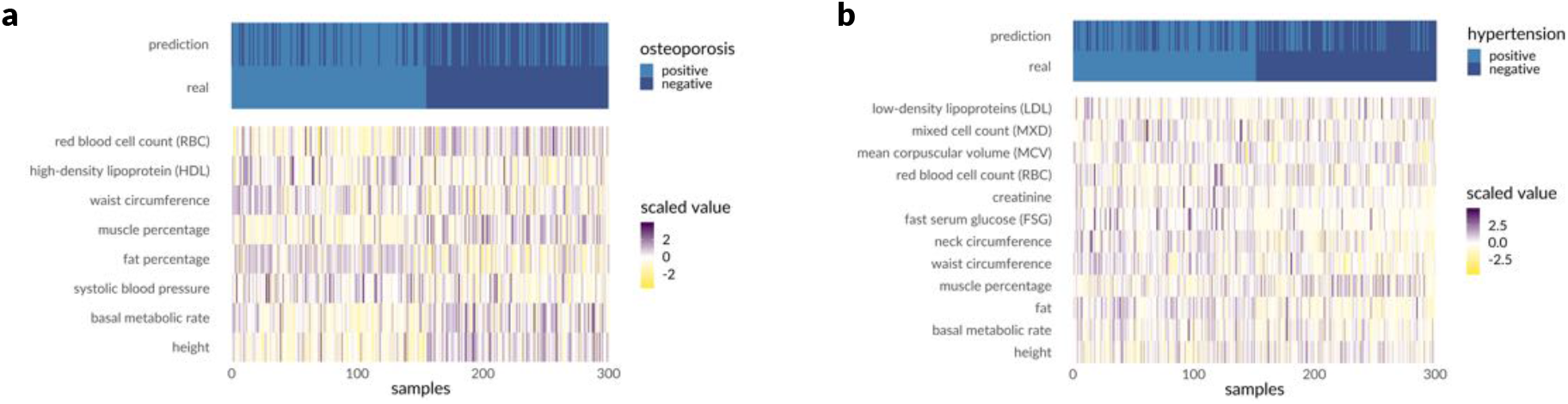
Multivariate elastic net model for categorical variables. **(a-b) An overview of the prediction model:** The predictors with a non-zero coefficient of in the model are taken into account. The scaled values of the predictors for a subsample of 100 participants are shown in the heat map. A heat map depicting the real values as well as the predicted values of the response variable is illustrated on the top, where the samples are sorted for the value of the response variable. **(a)** Osteoporosis, **(b)** hypertension.

## Discussion (in progress)

Health surveys are one of the most informative resources about the populations. By measuring various features from a subpopulation in a health survey, researchers may learn considerably about the inherent characteristics of the population under study. Such knowledge about the populations will be enhanced by increasing the sample size of surveys. From another perspective, surveys can be employed so as to find the relations among different features of a human population. Specifically, one of the most prominent discoveries is finding the biomarkers of a disease or predicting a measurement from other measurements. Although there has been a lot of health surveys performed, all over the world, there is an immense gap between what potentially can be elicited from data and contemporary knowledge. Such a gap can be addressed merely through rigorous quantitative methodologies. To date, there are numerous approaches proposed to the problem in hand; however, many of them are biased towards a limited set of hypotheses which may lead to confounding results due to neglecting the global structure of interactions among variables. Thus, tools capable of extracting information from health surveys in an unbiased manner is crucial.

In this article, we proposed an end-to-end analysis work flow which is assumed to answer commonly-asked questions in health survey analyses and provide an unbiased framework aimed to find the whole interactome among features at once. While the alignment of the majority of our findings with the previous medical knowledge demonstrates the performance of our work flow, the new findings are surprising and raise some hypotheses which worth follow-up experiments and analyses.

Our GGM can be precisely decomposed into four functional modules each of which can be labeled with special machinery. This persuades one that the GGM is accurately constructed. In the small-scale one can investigate single edges, as what we have illustrated in the supplementary material using scatter plots of two continuous variables. The GGM highlights the importance of neck circumference as a rather newly-proposed body measurement. Neck circumference is the center of the GGM with the most degree in the graph. Moreover, this variable has the third most cumulative mutual information among the continuous variables.

The causal network suggests some causal relations between pairs of variables. Although many associations are estimated to be bi-directed, in line with the known natural feedback between some pairs, a few links are single directed and of high interest. Among those, the causal relationship between neck circumference and fat as well as the relationship between creatinine and uric acid (UAC) are very intriguing.

Our minimal forest perfectly captures the functional structure of the variables in high dimensions. Each graph community in the minimal forest shows to be a particular functional module. Furthermore, the single pair relations are completely accepted or acceptable by the medical community which indicates how effectively such models fit the problem of survey analysis. Albeit the intricate network of connections demands a comprehensive investigation on every single edge, a subset of results reported in this article is remarkable. The significant difference in creatinine levels between males and females, the relation between systolic blood pressure and history of kidney stone surgery, and the difference between HDL levels of males and females area among the results. Moreover, our minimal forest could efficiently detect the effect of familial history in asthma and AD on the vulnerability of these diseases. Our minimal forest also finds the difference between the tendency of thyroid problems between men and women.

Our proposed VKL well suits finding disease biomarkers, by detecting the variables which differ the most between affected and unaffected groups regarding distribution. In this article, we reported some of the results of our VKL analyses on heart diseases, thyroid problems, and osteoporosis — the findings of which can be introduced as candidate biomarkers of these diseases. Besides, our VVKL method surprisingly finds FSG as the feature which violates the linear relation between fat and systolic blood pressure.

The multivariate elastic net models could finely predict the response variables using continuous variables. The results are convincing about the efficacy of such models in prediction. With which one can use the formula of the model for further predictions.

All in all, our comprehensive analysis led to numerous findings each of which demands more focused statistical and experimental validations. This study has gone some way towards enhancing the methodological practice of health survey analyses. Further, The work flow may have many implications in survey analyses in other fields, such as social sciences, economics, and environmental research. Results so far have been very encouraging on the Iranian population, and the work flow should also be examined on the other populations.

Further experimental studies are needed to evaluate the proposed work flow. To that end, our colleagues in the Yazd cardiovascular research center are acquiring the second batch of the data on the same set of participants. This data would be used by us to perform a longitudinal study and validate our hypotheses and prediction models, as well. We are currently developing an **R** shiny application to provide the community with an interface to our work flow on the YaHS dataset.

**Table 1:** Details of the continuous variables: id, variable description, degree in GGM, betweenness centrality in GGM, summation of the mutual information with other variables.

| id | variable | degree | betweenness centrality | mutual information |
| --- | --- | --- | --- | --- |
| NeckCircum | neck circumference | 11 | 88.35 | 16.55 |
| BaseMeta | basal metabolic rate | 10 | 50.19 | 7.67 |
| Fat | fat | 9 | 42.73 | 17.27 |
| HB | hemoglobin | 9 | 127.45 | 14.78 |
| FatPerc | fat percentage | 8 | 74.82 | 7.96 |
| Weight | weight | 7 | 3.87 | 9.23 |
| MusclePerc | muscle percentage | 7 | 9.69 | 8.97 |
| Crea | creatinine | 7 | 21.94 | 19.22 |
| RBC | red blood cell count | 7 | 34.26 | 8.40 |
| Height | height | 6 | 7.11 | 15.54 |
| BMI | body mass index | 6 | 5.60 | 8.96 |
| WaistCircum | waist circumference | 6 | 6.43 | 10.92 |
| HipCircum | hip circumference | 6 | 4.40 | 12.91 |
| UAC | uric acid | 6 | 32.75 | 12.50 |
| MCH | mean cell hemoglobin | 6 | 31.95 | 12.91 |
| HDL | high-density lipoprotein | 5 | 37.86 | 12.93 |
| Platelet | platelet | 5 | 141.52 | 8.76 |
| TG | triglyceride | 4 | 19.02 | 14.35 |
| Hct | hematocrit | 4 | 13.31 | 10.30 |
| BloodPreS | systolic blood pressure | 3 | 15.52 | 10.59 |
| MCV | mean corpuscular volume | 3 | 1.51 | 13.85 |
| MCHC | mean corpuscular hemoglobin concentration | 3 | 6.68 | 13.77 |
| Neut | neutrophils | 3 | 58.00 | 9.54 |
| RDW | red blood cell distribution width | 3 | 2.33 | 12.10 |
| PDW | platelet distribution width | 3 | 5.08 | 13.78 |
| MPV | mean platelet volume | 3 | 6.64 | 13.45 |
| BloodPreD | diastolic blood pressure | 2 | 5.44 | 12.18 |
| FSG | fast serum glucose | 2 | 2.53 | 15.36 |
| Chol | cholesterol | 2 | 0.00 | 7.90 |
| WBC | white blood cell count | 2 | 84.00 | 10.66 |
| Lymph | lymph | 2 | 0.00 | 8.20 |
| Mxd | mixed cell count | 2 | 0.00 | 7.93 |
| Pulse | pulse | 0 | 0.00 | 11.95 |

## Acknowledgements

Authors would like to acknowledge the Yazd people for participation in the study, graduate students for participation in design and Yazd Central Health workers and managers for acquiring the data.

## Author Contributions

E. H. devised the methodological work flow. V. B. and E. H. implemented the work flow. E. H. wrote the manuscript. N. A, M. S., and N. A. performed the survey and provided the data. A. S., M. M., and M. S. revised the manuscript. N. A. directed the collaboration. M. M. is the chief investigator of Yazd Health Study.

## Online Methods

### 7.1 The YaHS dataset

Yazd Health Study (YaHS) is a population-based cohort study of 10000 individuals aged uniformly from 20 to 70 years, conducted in November 2014 in the Greater Yazd Area of Iran. A verified questionnaire including 300 questions regarding a) demographics, b) physical activity, c) sleep quality and quantity, d) mental health, e) past medical history of chronic disease and surgical operations, f) dental health, g) history of accidents, h) dietary habits, i) occupation and social life, j) traditional medicine, k) smoking habit and drug addiction, l) women’s health, and m) quality of life is recorded by trained interviewers as well as anthropometrics, blood pressure, and vital signs from 4010 participants.

### 7.2 Uniform Manifold Approximation and Projection (UMAP)

UMAP is a non-linear dimension reduction method. It can be used to approximate a manifold on which the data is assumed to lie and embed it in a low dimensional projection in which the embedding is the closest possible equivalent fuzzy topological structure [30]. We use UMAP to embed the population in 2 dimensions.

### 7.3 Variable-wise Kullback-Leibler divergence (VKL)

Kullback-Leibler divergence (KL-divergence) is a measure of the deviation of one probability distribution from one another. As explained in [27], KL-divergence can be summarized to be then used as a symmetric measure of dissimilarity between two probability distributions. Given two disjoint groups of samples, one can calculate KL-divergence for each variable in the dataset to see how different are the probability distributions of the variables between the two groups of samples. Such an approach is called variable-wise Kullback-Leibler divergence (VKL). We use R package **muvis** [31] to conduct VKL on the dataset. The advantage of using KL-divergence in our setting is the feasibility of its usage on both categorical (i.e., questions) and continuous variables (i.e., laboratory and body measurements).

Performing VKL on the dataset, for each variable one can get a value indicating how the variable is distinct between the two groups above of samples. Thus, this value can be interpreted to find the most important variables associated with the discrimination of the groups.

### 7.4 Violating variable-wise Kullback-Leibler divergence (VVKL)

VVKL generalizes VKL. The method fits a linear regression model on two variables and remark samples with distances above a predetermined cut-off from the fitted line as outliers. Those outliers are supposed to bear some features which violate the expected linear association between the two variables. The groups are recognized as samples above and under the fitted line, respectively. Those groups, after that, are passed to VKL method. The output of the VKL method can be used to find the variables which violate the aforesaid linear association. Again, we use **muvis** to perform VVKL.

### 7.5 Graphical representations

We depict the estimated graphical models as graphs. In those such graphs, each node represents a variable, and two nodes are connected by an (indirect or directed) edge if the variables are estimated to be related based on the graphical model. Nodes are colored based on their node community, and their size indicates their betweenness centrality. The graphs are visualized using **muvis** and **gephi** [32].

#### 7.5.1 Community detection algorithm

Investigating node communities within graphical models helps interpretation through highlighting highly-related sets of nodes which can be inferred as distinct functional modules of variables. To this end, we used the Lovian community detection algorithm [33]. Hence, each community can be recognized by a specific color in the graph.

#### 7.5.2 Community detection algorithm

Betweenness centrality is a measure of node centrality in a graph and is defined as the number of shortest paths between pairs of nodes which contain the node of interest. This measure, thus, can indicate how important is a node in a graph. Particularly in graphical models in which edges present statistical associations, betweenness centrality is a prominent criterion to show how a node contributes to relations among the variables.

### 7.6 Graphical models (GMs)

GMs are widely-known for delineation of relations among a set of variables. Based on observed data, GMs are aimed to estimate and represent the rather intricate structure of associations among variables, in an intelligible manner[34]. We use three different types of graphical models, that is, Gaussian graphical model (GGM), minimal forest, and causal network. To that end, we used the implementation of these graphical models in the **muvis** package.

#### 7.6.1 Gaussian graphical model (GGM)

glasso–graphical lasso, an algorithm to construct GGMs-is employed to estimate associations among continuous variables [35]. In the GGM each node represents a continuous variable and each edge shows non-zero partial correlation (i.e., the correlation between two variables given all other variables) between two variables. GGM, by its nature, leads to a sparse representation of the relations which is straightforward to interpret.

#### 7.6.2 Minimal forest

Considering both continuous and categorical variables, we exploit minimal forest to create a graphical model on the whole set of variables. Minimal forest uses BIC algorithm to build a sparse graphical model on a large set of variables [36]. In minimal forest, each node represents a (continuous or categorical) variable and each edge represents a penalized mutual information using BIC algorithm.

#### 7.6.3 Causal network

So as to find causal relations among continuous variables, that is to say which variable is a cause another, we use causal network introduced in [37]. In the causal network, each node represents a continuous variable, and each (directed) edge indicates a causal relationship. The direction of edges depicts the direction of causal relation, from cause to effect.

### 7.7 The Elastic net model

The Elastic net is a regularized linear regression or logistic regression [38]. The elastic net regularization is a linear combination of l1 and l2 penalization of the lasso (least absolute shrinkage and selection operator) and the ridge regressions. Manipulating the hyperparameter of elastic net penalty one can enjoy the advantages of both lasso and ridge regression. Furthermore, it has been shown that elastic net selects or excludes groups of variables which are highly correlated [38] — these characteristics of the elastic net benefits one to find functional modules that contribute to the prediction of a variable. Thus, we use the elastic net model form **muvis** to find the variables which are important to predict a variable thereby finding the most important associations with a variable of interest. The same package is used to visualize the learned elastic net model.

